# Solid-state fermentation with mushroom mycelium elevates plant protein quality and umami taste

**DOI:** 10.1101/2025.09.24.678313

**Authors:** Jasper Zwinkels, Eddy J. Smid, Oscar van Mastrigt

## Abstract

Solid-state fungal fermentation (SSFF) offers a low-tech, low-energy, and minimal processing method to enhance the protein content and quality of foods. This study evaluates the potential of SSFF executed with mycelium of edible basidiomycetes to improve both nutritional and sensory qualities of brown rice, brewer’s spent grain (BSG), and lupin. The conventional tempeh fungus, *Rhizopus microsporus* var. *oligosporus*, was used as control. Different substrate-fungus combinations varied in impact on flavour and protein quality. SSFF improved the protein quality and umami taste of brown rice and mainly improved in umami taste of lupin, while fermentation of BSG with basidiomycetes even decreased protein quality. Basidiomycetous SSFF products exhibited higher umami potential, with equivalent umami concentrations (EUC) reaching 159 g MSG-eq/100 g DW, surpassing the values found for *R. microsporus*-fermented products. In terms of substrates, the protein content increased most in brown rice fermentations, while the EUC and protein quality increased most in lupin. Protein quality and content increased more in basidiomycetes, indicated by the up to 35.1% increase in the protein digestibility corrected amino acid score of the limiting amino acid lysine and a 36.4% rise in utilizable amino acids. Basidiomycetous SSFF thus offers a promising approach to upgrade low-quality plant proteins into more palatable and nutritious foods.

**Highlights:**

– Plant foods were fermented with basidiomycetes and conventional tempeh fungus. (81)
– Basidiomycetes improve umami compounds more than conventional fermentation fungus. (85)
– Basidiomycetes improve protein quality more than conventional food fungus. (84)
– Increase in total utilizable amino acids requires proper substrate-fungus pairing. (85)
– Basidiomycota harbours untapped phylogenetic diversity for food fermentation. (79)

## 1 Introduction

In developed nations a higher portion of the protein intake needs to come from non-animal proteins for sustainability, health, and ethical reasons. Higher consumption of (red) meat is associated with i.a. higher cardiovascular disease incidence, higher greenhouse gas emissions, and deforestation (Kim et al., 2019; Poore & Nemecek, 2018). Despite the fact that median protein intakes generally exceed requirements in developed nations, vegan and vegetarian diets lead to a higher incidence of inadequate protein intakes (Broekema et al., 2020; Kim et al., 2019). Additionally, in developing nations a substantial portion of the protein intake comes from plant sources, like cereals and roots. The staple food group cereals contribute to 30% of the total protein intake (FAO, 2024). These staple foods are characterized by a low protein quality (Sá & House, 2024), and this, and this leads to an increase in the prevalence of protein inadequacy (Moughan, 2021). Therefore, in developing as well as in developed nations, a higher proportion of high-quality plant protein in the diet could alleviate these issues.

A method to increase the protein quality of cereals and legumes is solid-state fungal fermentation (SSFF) (Villacrés & Rosell, 2021; Zwinkels et al., 2023). This minimal processing method can improve the digestibility, as well as increase the amount of utilizable lysine, which is commonly the limiting amino acid in these foods. SSFF has a particular advantage over other processing methods as it minimally impacts other beneficial nutrients, such as fibres and minerals, and does not require the addition of salt or other preservatives. Moreover, low-income populations often have little access to healthy and sustainable protein alternatives (Lumsden et al., 2024). The low energy and technological requirements of SSFF make it a technique that can be widely applied in many parts of the world. Currently, SSFF relies mainly on a limited set of fungi: mucoromycetes such as *Rhizopus* sp. (tempeh), and ascomycetes including *Aspergillus* sp. (koji), *Neurospora intermedia* (oncom), and *Penicillium* sp. (mould-ripened cheeses) (Allwood et al., 2021; Finnigan, 2011; Han et al., 2004; Starzynska-Janiszewska et al., 2017; Wolkers – Rooijackers et al., 2018; Yin et al., 2020).

Besides Mucoromycota and Ascomycota, the fungal kingdom harbours one more phylum that has potential, particularly for food fermentation. The phylum Basidiomycota is composed of roughly 40,000 species, of which around 2,000 produce edible fruiting bodies (mushrooms) (He et al., 2022).

These mushrooms have a long history of consumption and are desired for their umami taste. This makes them interesting options for meat alternatives (Mau, 2005; Miller et al., 2014) as umami taste is a key factor in alternative protein acceptability and sustainable eating (Schmidt & Mouritsen, 2022; Yamaguchi & Ninomiya, 2000). However, these mushrooms generally have a lower protein quality and content, making them less ideal to meet dietary protein requirements (Wallis et al., 2012; Zwinkels, van Oorschot, et al., 2025). Nonetheless, the mycelium of these species can still be a good potential source of dietary protein. A recent study indicated that the mycelium of basidiomycetes possesses both a high content of umami-active compounds, as well as a high protein content and quality, outperforming the conventional fungi used in solid-state fermentation for tempeh production (Zwinkels, van Oorschot, et al., 2025). Secondly, basidiomycetes are well-known to grow on lignocellulosic material, commonly found in food industry side-streams, making them a potentially sustainable solution for protein production (Ilić et al., 2023). Thirdly, the mycelium of the basidiomycete *Pleurotus ostreatus* did not contain known mycotoxins. Moreover, four peptide toxins were lower in mycelium compared to the fruiting body (van Dam et al., 2024). A strong indication that mycelium of edible mushroom-producing fungi is safe for human consumption. Lastly, the mycelium of five basidiomycete species was found to grow on plant substrates suitable for human consumption, where it reduced the anti-nutritional factor phytic acid more effectively than *R. microsporus* (Zwinkels, Oorschot, et al., 2025). The four above-mentioned factors support the potential of basidiomycetous mycelium in SSFF as a healthy, sustainable, and tasty novel protein source.

However to date, research on basidiomycetes has predominantly focused on mushroom production, enzyme production, waste-stream valorisation for feed production, or pure biomass production through submerged fermentation (Ahlborn et al., 2019; Bach et al., 2017; da Silveira & Badiale-Furlong, 2009; Henske et al., 2018; Stephan et al., 2018; Suruga et al., 2020; van Dam et al., 2024).In contrast, research on SSFF of food-grade substrates with basidiomycetous mycelia and their effect on protein quality and flavour is scarce.

Therefore, this study aims to evaluate the potential of SSFF foods produced with basidiomycetous mycelia as novel protein sources for human nutrition by assessing their protein content and quality (via protein digestibility corrected amino acid score (PDCAAS)), alongside the presence of umami-active molecules, and roasted meat-like aromas. Tempeh made with *Rhizopus microsporus* var. *oligosporus* on the same substrates serves as a reference.

To produce a protein source fit for human consumption through SSFF, without requiring additional processing techniques, plant substrates fit for human consumption are required. Therefore, three substrates are selected that represent food groups used in food fermentation. Firstly, brown rice is a cereal and staple food in many parts of the world. It is high in starch and low in protein, but due to its high consumption contributing to a substantial portion of dietary protein intake in some areas (FAO, 2023). Lupin is a legume with a high protein and fibre content (Martínez-Villaluenga et al., 2006). Furthermore, it serves as a sustainable alternative to soy, due to its nitrogen fixation and possibility of cultivation in temperate regions (Palmero et al., 2022). Lastly, brewer’s spent grain (BSG) is an industry side-stream of beer brewing, made from malted barley. It has a low starch content, due to the malting and mashing process and is high in fibre and moderately high in protein (Briggs et al., 1981).

## 2 Materials & Methods

### 2.1 Substrates, species, and solid-state fermentation

Substrates and fungal species used, as well as the fermentation process are described in Zwinkels, Oorschot, et al. (2025). In brief, basmati brown rice (*Oryza sativa*), brewer’s spent grain (BSG, *Hordeum vulgare*), and white lupin (*Lupinus albus*) were fermented with the basidiomycetes *Stropharia rugoso-annulata, Schizophyllum commune, Volvariella volvacea, Pleurotus pulmonarius*, and *Pycnoporus cinnabarinus*. For comparison to conventionally used fungi for fermentation, these substrates were also fermented with *Rhizopus microsporus* var. *oligosporus*, commonly used in tempeh production.

Substrates were soaked overnight at pH 4.0, drained, rinsed, and autoclaved in a Microbox polypropylene fermentation container with an air filter strip. The substrates were inoculated with a mycelium suspension in the case of basidiomycetes or a spore suspension in the case of *R. microsporus*. Containers were incubated at the optimal growth temperature of the fungi (determined in Zwinkels, Oorschot, et al. (2025), which was 33°C for *P. cinnabarinus* and *V. volvacea*, 30°C for *P. pulmonarius* and *S. commune*, 27°C for *S. rugoso-annulata* and 28°C for *R. microsporus*. Fermentation time for basidiomycetes was eight days and for *R. microsporus* was 48 hours.

### 2.2 Sample preparation

After fermentation, the fermented cake was cut into 8 identical slices and pieces from all sides of the sample were taken, frozen with liquid nitrogen and ground to a fine powder using a spice grinder (Krups F203, Solingen, Germany). Homogenized samples were immediately stored in a urine container (VWR international, ML15) at −20°C until further analysis. Dry matter content was calculated from samples (1.0 g) dried to a constant weight in a moisture analyser MA100 (Sartorius, Germany) at 105°C, following the procedures described by the Association of Official Agricultural Chemists (AOAC, 2005).

### 2.3 Quantification of umami-active and volatile metabolites

#### 2.3.1 Free amino acids

Free amino acid concentrations in homogenized samples were determined by following the method described by Scott et al. (2021).

### 2.3.2 ‘5-nucleotides

5’-nucleotides were extracted and measured according to the method from Seifar et al. (2009). Homogenized samples (100 mg) were dispersed in 5 mL milliQ water in a 15 mL centrifugal tube. Tubes were mixed by vortexing and heated for 2 min at 100°C in a shaking water bath. The heated tubes were vortexed for 30 seconds and cooled to room temperature. Samples were centrifuged (5 min, 10.000 g) and the supernatant was filtered using a Costar Spin-X centrifuge filter (0.22 µm, cellulose acetate). Subsequently, 200 µL was transferred to a plastic autosampler vial, capped and stored at −20°C until analysis. Separation, detection and quantification of 5’-nucleotides was performed using ultraperformance liquid chromatography (UPLC) with an UltiMate 3000 UPLC system (Dionex, Idstein, Germany) equipped with an AccQ-Tag Ultra BEH C_18_ column (150 mm x 2.1 mm, 1.7 µm) (Waters, Milford, MA, USA), and a BEH C_18_ guard column (5 mm x 2.1 mm, 1.7 µm; Waters, Milford, MA, USA). The column temperature was set 25°C and the mobile phase flow rate was maintained at 0.2 ml/min. Eluent A was composed of 10 mM dibutyl ammonium acetate (DBAA, pH 6.7) (TCI chemicals, Tokyo, Japan) and eluent B was composed of 10 mM DBAA (pH 6.7) containing 84% acetonitrile (Supelco, Bellefonte, PA, USA). The separation gradient was 0 – 2 min 99.9% A, 2 – 12 min steadily decreasing to 70% A, 12 – 15 min 70% A, 15 – 17 min 0% A, 17 – 27 min 99.9% A. One microliter of sample was injected for analysis and compounds were detected by UV measurements at 250 nm. Concentrations were calculated based on a calibration curve of 5’-adenosine monophosphate (Thermo Fisher), 5’-guanosine monophosphate (Thermo Fisher), 5’-inosine monophosphate (Thermo Fisher), and 5’-xanthosine monophosphate (Bio-Connect, Huissen, The Netherlands) (0.78 – 25 mg/L).

#### 2.3.3 Equivalent umami concentration

The equivalent umami concentration (EUC) expresses the synergistic effect of umami free amino acid and 5’-nucleotides. Based on the relative umami concentration of these compounds, the EUC of MSG-eq perceived was calculated (Equation 1;Yamaguchi et al., 1971).

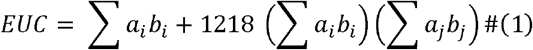

Where *a_i_* (g/100 g DW) is the concentration of the respective umami amino acid (Asp or Glu); *a*_j_ (g/100 g DW) is the concentration of the respective umami 5’-nucleotide (5⍰-MP, 5⍰-GMP, 5⍰ -XMP, or 5⍰ -AMP); *b*_i_ is the relative umami concentration (RUC) for umami amino acids to MSG; and *b*_j_ is the RUC for umami 5⍰ -nucleotides to 5⍰ -IMP (Yamaguchi et al., 1971).

#### 2.3.4 Volatile organic compounds

Volatile organic compounds (VOCs), with a focus on alkylpyrazines, were determined by headspace solid-phase microextraction gas chromatography mass spectrometry (HS SPME GC-MS) (Scott et al., 2021). In brief, 1.0 gram homogenized samples were weighed in a 5 mL gas chromatography vial. Raw samples were capped and stored at −20°C until analysis. Heated samples were covered with aluminium foil and heated in a convection oven at 150°C for 5 min, after which they were also capped and stored at −20°C. VOCs were extracted and detected according to the method described by Scott et al. (2021), with the modification that a 1:5 split mode was used.

### 2.4 Protein quality

#### 2.4.1 *In vitro* protein digestibility

*In vitro* protein digestibility (IVPD) was determined using the Megazyme test kit K-PDCAAS by following the manufacturer’s procedure with some adaptations (Megazyme, 2019). In brief, approximately 500 mg homogenized test, blank, control and standard sample was weighed into a 50 mL centrifugal tube. To these tubes 19 mL HCl (0.06 M) was added, and the tubes were incubated in a hot air shaking incubator (37°C, 30 min, 300 RPM). This was followed by a first digestion step with the addition of pepsin (60 min, 37°C, 300 RPM). Subsequently, samples were neutralized to pH 7.4 by addition of 2 mL 1.0 M Tris buffer. Then, samples were subjected to a second digestion step by addition of 200 µL trypsin/chymotrypsin mixture and incubation for 4 h at 37°C at 300 RPM. This was followed by heating the samples in a boiling water bath for 10 min. Proteins were precipitated overnight at 4°C by addition of 1 mL TCA solution (40%). Tubes were centrifuged (10 min, 15,000 g) and the supernatants were diluted 10-fold and 20-fold in acetate buffer (50 mM, pH 5.5). Amine concentration was measured against a L-glycine calibration curve through colorimetric determination of test, blank and standard samples at 570 nm after the addition of ninhydrin (2%) and incubation at 70°C for 35 min. IVPD was calculated from the primary amine concentration of test samples fitted against standard samples with known *in vivo* protein digestibility values.

#### 2.4.2 Amino acid composition

The processing, extraction, and quantification of amino acids in the samples are described in Zwinkels, Oorschot, et al. (2025). The procedures guided by ISO13904 (ISO, 2005b) and ISO13903 (ISO, 2005a) were followed. Lupin products were defatted before hydrolysis.

#### 2.4.3 Protein quality and utilizable amino acid content

Protein quality was determined based on the protein digestibility corrected amino acid score (PDCAAS), as described by (FAO/WHO, 1991). Some adaptations were made to make the score more in line with the digestible indispensable amino acid score (DIAAS), which was recommended by the FAO (2013). The adaptations were: (1) the inclusion of non-limiting amino acids in the score, and (2) not truncating scores above 1.0. The amino acid score (AAS) was calculated using Equation 2. The reference protein was based on the dietary reference intake of a pre-school child (0.5–3 y), as prescribed by the FAO (2013).

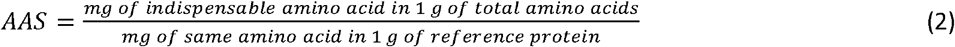

The PDCAAS was calculated by multiplying the IVPD by the AAS of the limiting amino acid (Equation 3)

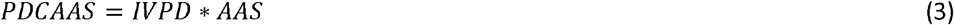

The utilizable amino acid content (UAA) was calculated by multiplying the truncated PDCAAS with the total amino acid content.

### 2.5 Data analysis

Data generated by UPLC and HS SPME GC-MS were analysed in Chromeleon 7.3.1 (Thermo Scientific, MA, USA). GC-MS peak integration was performed using the ICIS algorithm and the NIST main library was used for identification by matching mass spectral profiles with the profiles in NIST. One quantifying peak (generally the highest *m/z* peak per compound) was used per compound for quantification, while two confirming peaks were used for confirmation. Data was analysed, and figures were produced using Rstudio software v. 4.0.2 (RStudio®, Boston, MA, USA). All values presented are means of biological triplicates ± standard deviation, unless stated otherwise. Significant differences that are indicated with letters were determined using One-Way ANOVA with Tukey post hoc test. Significance level (*p*-value) was set at 0.05.

## 3 Results

### 3.1 Taste & aroma

The effect of solid-sate fermentation by novel basidiomycetous fungi on umami taste compounds of cereals and legumes was compared to the conventional food fungus *R. microsporus*, used in tempeh production. The equivalent umami concentration (EUC) was calculated as a measure for umami taste of the food product (**Figure 1A**). The unfermented substrates, BSG, brown rice, and lupin, had a low EUC of 0.007, 0.028 and 1.52 g MSG-eq/100 g DW, respectively, placing them in umami level 4 (0-10 g MSG-eq/100 g DW), according to Mau (2005). Fermentation increased the EUC in all samples, but the extend of this increase was dependent on the interaction between substrate and fungal species. EUC increased to level 3 (10-100 g MSG-eq/100 g DW) in *P. pulmonarius* fermented BSG and brown rice, *R. microsporus*-lupin and *P. cinnabarinus*-brown rice. The highest EUC was obtained in *P. cinnabarinus*-lupin (158.7 g MSG-eq/100 g DW), reaching level 2. Generally, the highest EUC was observed in lupin, then brown rice, followed by BSG.

**Figure 1.**
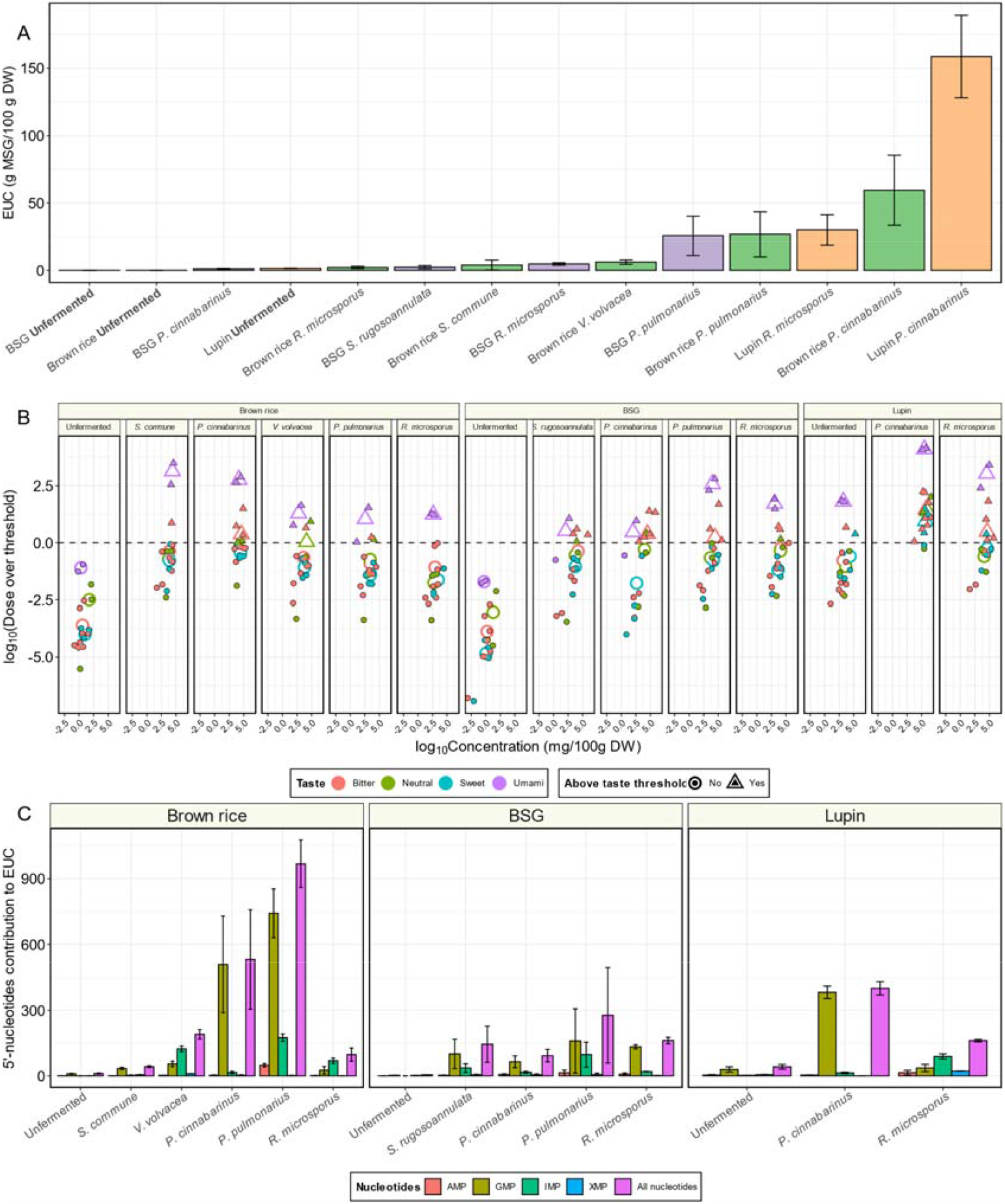
**A)** Equivalent umami concentration (EUC, g MSG-eq/100 g DW) of unfermented and fermented lupin (orange), brown rice (green) and brewer’s spent grain (BSG, purple) (mean ± standard deviation). **B)** Relationship between log_10_(Dose over threshold) and log_10_(concentration) of free amino acids with bitter (red), neutral (green), sweet (blue), or umami (purple) taste in unfermented and fermented substrates. Closed points represent individual amino acids, while open points represent the average of all amino acids in a certain taste category. Round points are below the taste threshold, while triangular points are above the taste threshold. **C)** Contribution of 5’-nucleotides to the equivalent umami concentration of unfermented and fermented substrates (Equation 1). Colours represent 5’-adenosine monophosphate (AMP, red), 5’-guanosine monophosphate (GMP, olive green), 5’-inosine monophosphate (IMP, green), 5’-xanthanosine monophosphate (XMP, blue), and the sum of all nucleotides contributions (purple).

**Figure 2.**
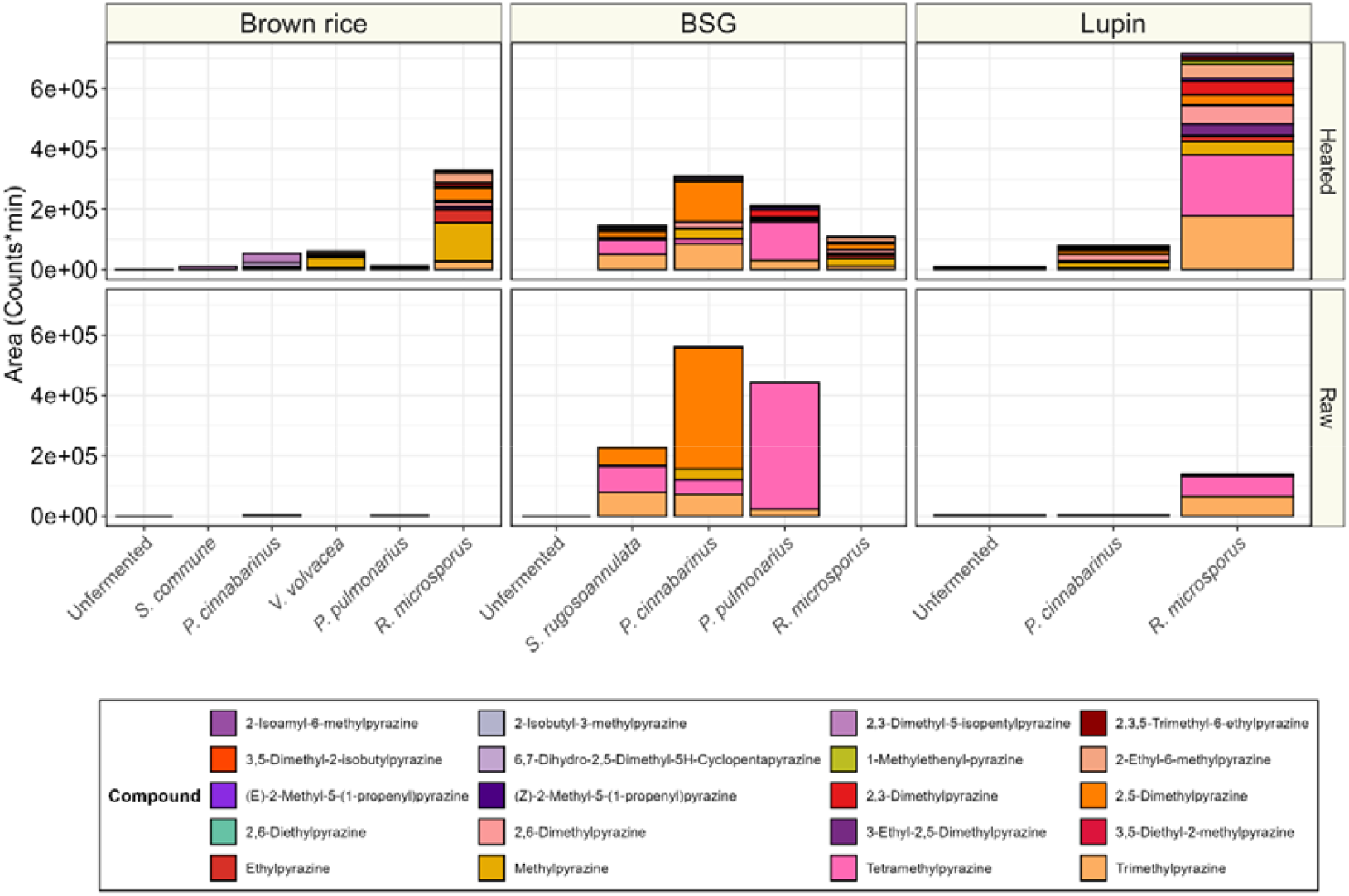
Alkylpyrazine areas (counts*min) per gram of dry weight unfermented or fermented substrate. Samples were measured either untreated (Raw) or after heat treatment (Heated). Colours represent compounds and are explained in the figure legend.

The umami taste in food is derived from a synergistic effect of umami free amino acids (FAA) and 5’-nucleotides. In unfermented substrates the concentration of umami FAA was slightly lower than neutral, bitter, or sweet tasting FAA (**Figure 1B**). However, the concentrations of all FAA were insufficient to pass the taste threshold. Fermentation increased the concentration of FAA in all instances. However, mainly umami FAA surpassed the taste threshold, reaching dose-over-threshold (DOT) up to 60 in *P. cinnabarinus*-lupin. Umami FAAs exhibited a higher DOT compared to other tasting FAA, primarily due to their low taste threshold. Lastly, the contribution of 5’-nucleotides to the EUC is depicted in Figure 1C. Similarly to FAA, 5’-nucleotides were only present in minor amounts in the unfermented products. In most fermentations, 5’-guanosine monophosphate (GMP) contributed most to the EUC of the fermented products, except for *V. volvacea* fermented brown rice and *R. microsporus* fermented lupin, where 5’-inosine monophosphate (IMP) was the highest contributor. Overall, production of 5’-nucleotides was higher in basidiomycetous fermentations than in *R. microsporus*. Particularly, *P. cinnabarinus* and *P. pulmonarius* produced these compounds in considerable amounts. Basidiomycetes seemed to favour brown rice for 5’-nucleotide production, while *R. microsporus* produced more in lupin.

Alkylpyrazines are heterocyclic nitrogenous aromatic compounds, and their presence is associated with meaty and roasted aromas (Ren et al., 2024). Levels of these compounds in unfermented and fermented substrates, either raw or after heat treatment were determined by HS SPME GC-MS. Alkylpyrazines were only detected at trace levels in unfermented substrates. Fermentation with basidiomycetes particularly produced alkylpyrazines in BSG (up to 5.6*10^5^ counts*min) and in lupin fermented with *R. microsporus* (6.9*10^5^ counts*min). The most abundant compound was either 2,5-dimethylpyrazine or tetramethylpyrazine. Heating increased the alkylpyrazine concentration in *R. microsporus* fermented substrates (up to 7.0*10^5^ counts*min), while changes were less pronounced in basidiomycetes fermentations. The diversity of pyrazine-derivatives did increase with heating, from maximally five in the raw substrate (in *P. cinnabarinus* fermented BSG) to twelve different compounds in the heated substrates (in *R. microsporus* fermented lupin).

### 3.2 Protein quality

In addition to taste and aroma, the effect of fermentation by basidiomycetes on nutritional quality of the substrates was assessed. Protein quality was determined based on the *in vitro* protein digestibility and amino acid composition of the fermented products. (**Figure 3**). In unfermented brown rice, the protein digestibility was 82.3% and fermentation altered the protein digestibility with less than 1.7 percentage points. However, in BSG, fermentation significantly reduced the protein digestibility from 88.1%, to as low as 80.0% (*p* < 0.05), whereas in lupin fermented with *P. cinnabarinus* digestibility significantly increased with 7.5 percentage points, reaching 99.4% (*p* < 0.05). Overall, the effect of fermentation on protein digestibility was based on the interaction between substrate and fungal species. This was illustrated by the effect of *P. cinnabarinus* on protein digestibility, which did not alter in brown rice, reduced the digestibility in BSG, and increased digestibility in lupin.

**Figure 3.**
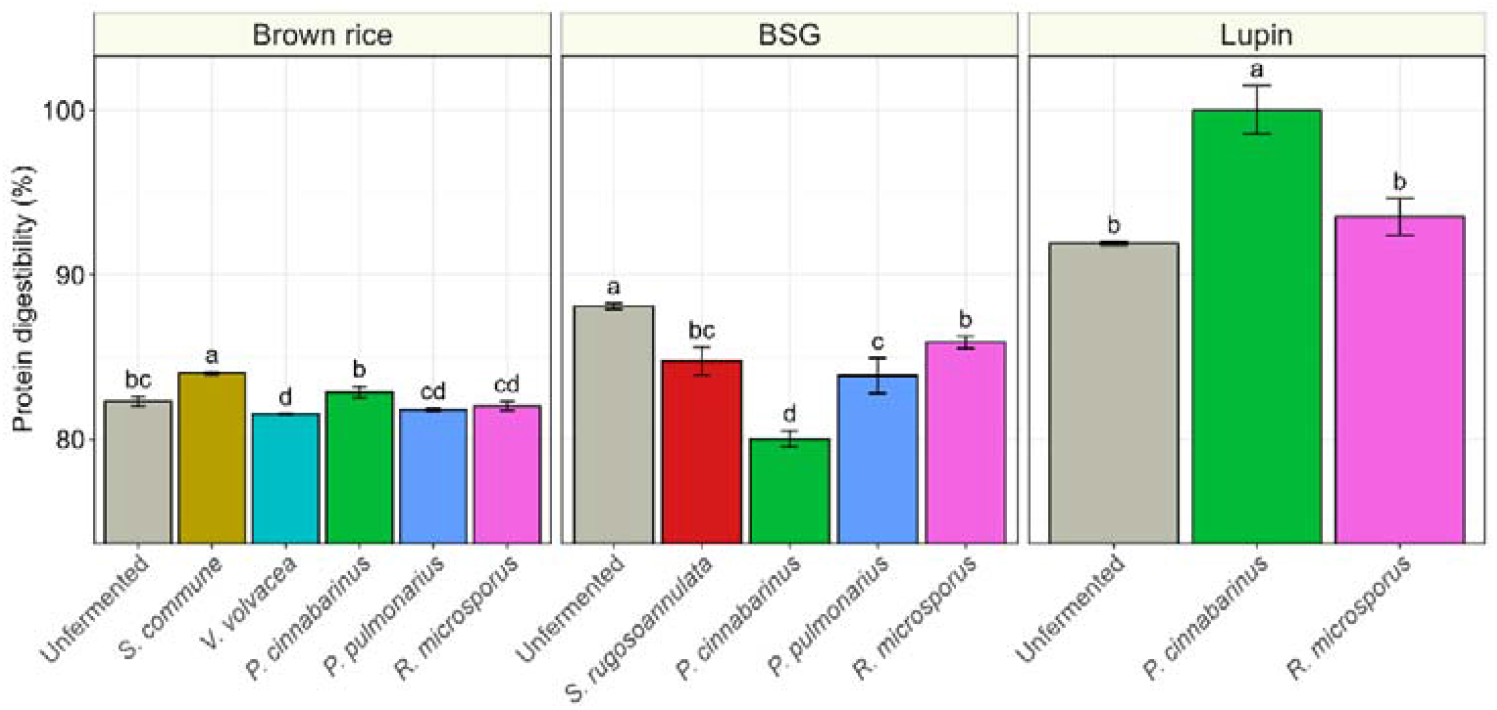
*In vitro* protein digestibility (%) of unfermented and fermented substrates. Different letters indicate significant differences between species within a substrate (TukeyHSD, *p* < 0.05).

Protein quality in unfermented and fermented substrates was determined based on the protein digestibility corrected amino acid score (PDCAAS) of all indispensable amino acids (**Figure 4**). In unfermented brown rice, lysine was the limiting amino acid, with a PDCAAS of 0.62. Fermentation increased the PDCAAS between 2.6 and 18.6%, up to 0.73. In addition to lysine, the PDCAAS similarly increased consistently in isoleucine and threonine. Furthermore, in BSG, lysine was the limiting amino acid with a PDCAAS of 0.78. Fermentation of BSG altered the PDCAAS depending on the fungal species. Fermentation with *P. pulmonarius* and *R. microsporus* had relatively minor effect on the PDCAAS, while *S. rugoso-annulata* fermentation increased the PDCAAS of lysine by 14.8% to 0.90. Fermentation with *P. cinnabarinus* substantially increased and decreased the protein digestibility corrected amino acid ratio (PDCAAR) of certain indispensable amino acids. On the other hand, lupin, in its unfermented state, was limiting in sulphur amino acids, lysine, and tryptophan, with a PDCAAS of tryptophan at 0.86. Fermentation improved PDCAAS. Particularly, in *P. cinnabarinus* fermented lupin, the PDCAAR of sulphur amino acids increased by 35.1%, lysine by 17.8%, and tryptophan by 52.7%. This fermentation increased the PDCAAS by such an extent that it can be considered a complete protein, with a PDCAAS of 1.03 (lysine). Similar to the increases in PDCAAS, fermentation significantly raised the proportion of indispensable amino acids across all substrates by 0.2–3.4% (Supplementary Figure 1), indicating that mycelium provides a more balanced amino acid composition than plants.

**Figure 4.**
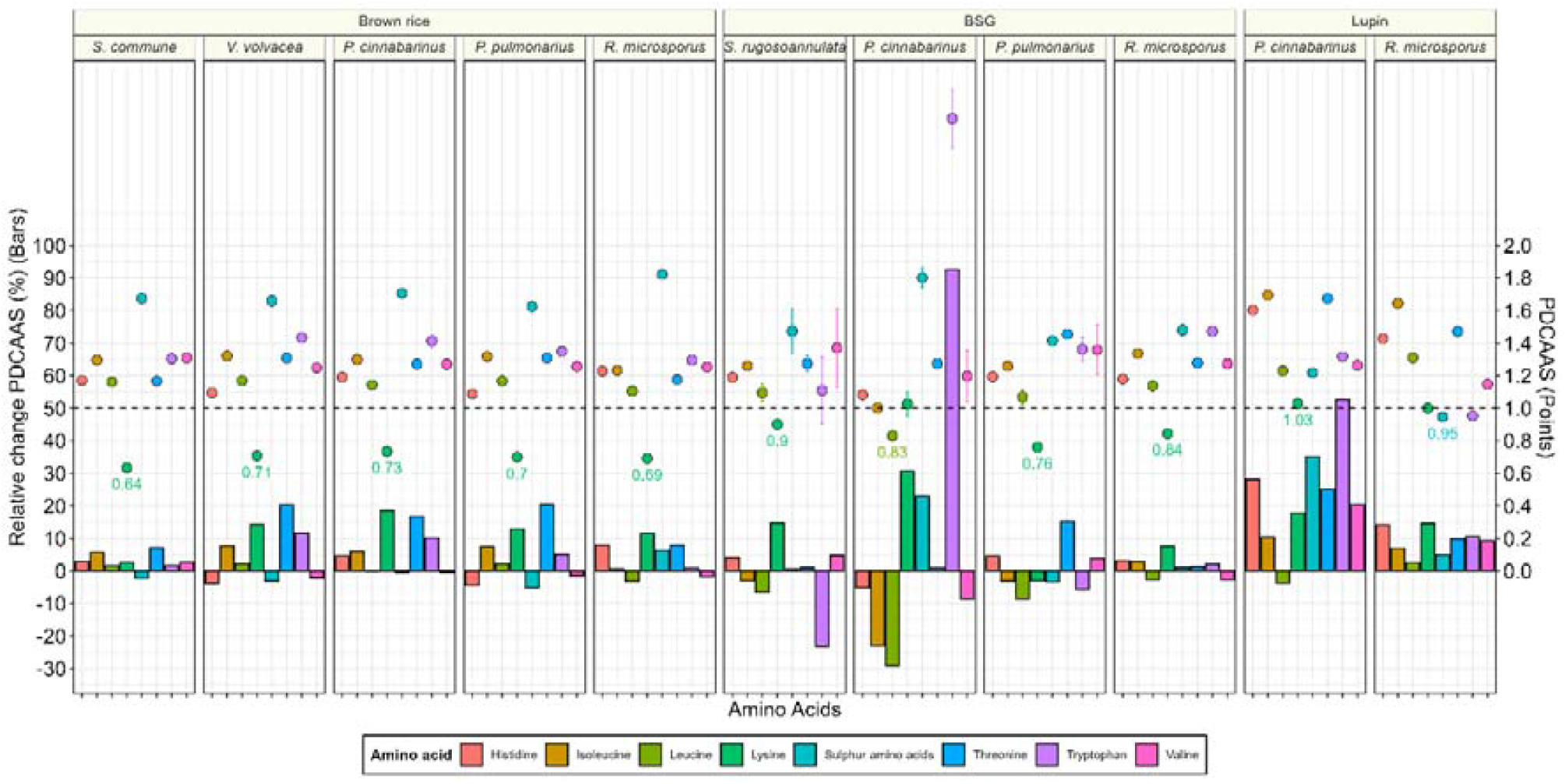
Protein quality of fermented substrates. Protein quality is represented by the protein digestibility corrected amino acid score (PDCAAS) of indispensable amino acids (coloured). Points represent the PDCAAS per amino acid (± standard deviation), while bars represent the relative change in PDCAAS from the unfermented substrate (%). The PDCAAS of the amino acids in the unfermented substrate is given in supplementary figure 2 and written in text for the limiting amino acid. Aromatic amino acids (phenylalanine and tyrosine) were not measured.

The increased PDCAAS of fermented substrates indicates that fungal fermentation enhances their protein quality. However, as described by Zwinkels, Oorschot, et al. (2025), fungal fermentation can also alter the protein content of the substrate through the breakdown of amino acids via deamination (decrease), or through loss of dry weight (increase). This was included through the calculation of utilizable amino acid content (UAA), indicating the total amino acid content that can be absorbed and utilized by the body. The change in UAA due to fermentation was plotted against the EUC (**Figure 5**). The UAA of unfermented brown rice, BSG, and lupin were 5.7, 14.6, and 31.2 g/100 g DW. In brown rice, the UAA increased by 15.3 to 36.4%, due to the combined increase in PDCAAS and increase in total amino acids content (Zwinkels, Oorschot, et al., 2025). Contrastingly, in BSG the change in UAA varied from 6.1% increase to 46.0% decrease, due to the degradation of amino acids, particularly present in BSG fermented with *P. cinnabarinus* and *P. pulmonarius*. In lupin, fermentation increased the UAA up to 6.1%. This increase in UAA was accompanied by the largest increase in EUC to 158.7 g MSG-eq/100 g DW.

**Figure 5.**
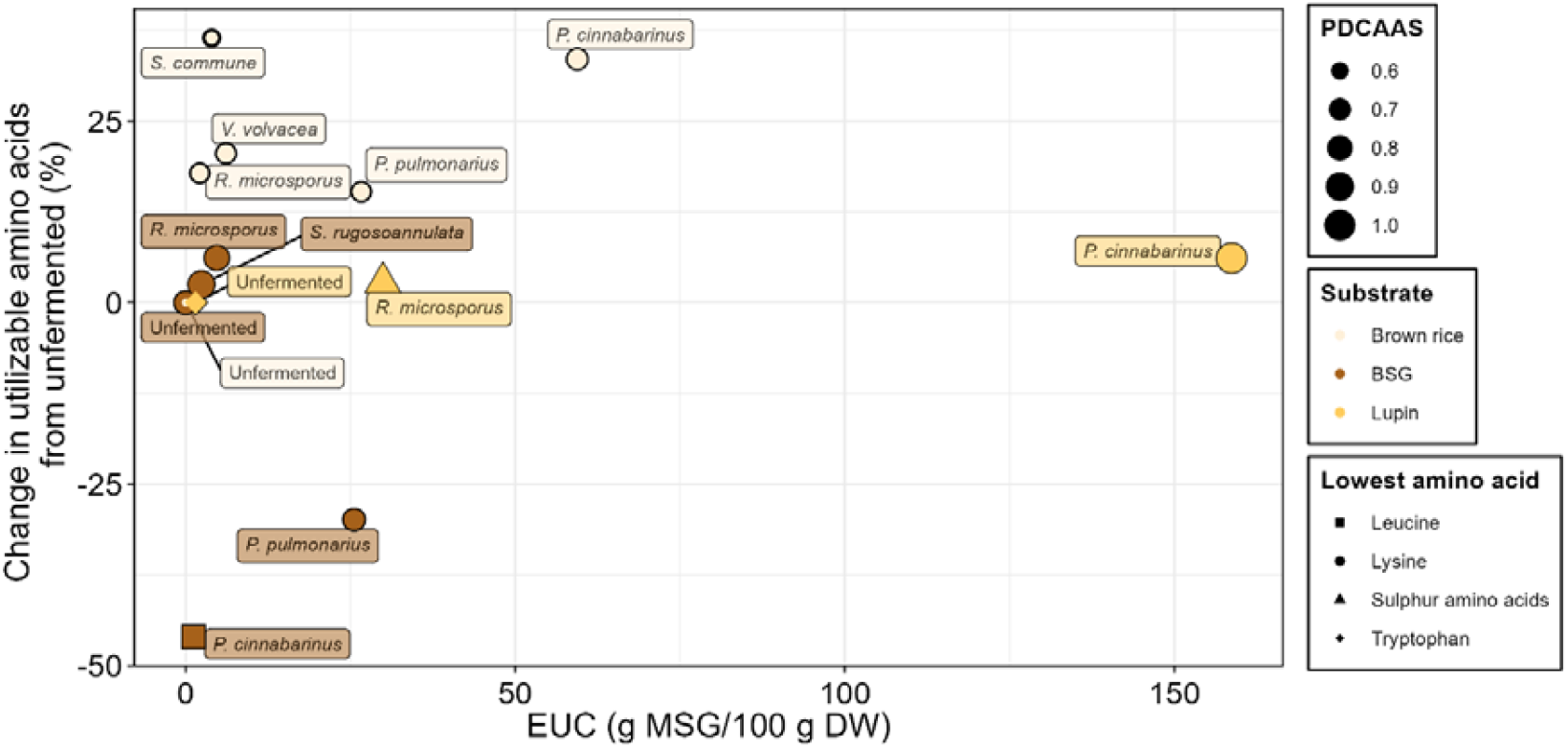
Effect of fermentation on the utilizable amino acid content (%) and equivalent umami concentration (EUC, g MSG-eq/100 g DW). To compare substrates, the change in the utilizable amino acid content compared to the unfermented substrate is given. PDCAAS of the sample is indicated by the size of the point. Colour of the point indicates the substrate. The shape of the point indicates the amino acid with the lower PDCAAS. Aromatic amino acids (phenylalanine and tyrosine) were not measured.

## 4 Discussion

This study explored the impact of solid-state fermentation with basidiomycetes on the taste, aroma, and protein quality of plant-based foods. The impact of basidiomycetes was also compared to fermentation with *R. microsporus*, representing a fungus that is conventionally used for solid-state food fermentation (i.e. tempeh making). With these assessments we aimed to understand the potential of solid-state fermentation with mycelium of basidiomycetes to produce novel high-quality protein sources for human consumption.

Umami taste is a critical factor in food likeability and novel food acceptance, with umami being called a major driver of sustainable food consumption (Schmidt & Mouritsen, 2022). The equivalent umami concentration (EUC), based on the synergistic effects of umami FAA and 5’-nucleotides, closely correlates with umami taste perception (Zhu et al., 2022). While meat is a well-known source of umami, mushrooms also have a particularly strong umami taste (Mau, 2005).

In our study basidiomycetes produced higher EUC values than *R. microsporus* across all substrates, reaching up to 159 g MSG-eq/100 g DW. The substantial EUC values in basidiomycetous SSFF products is in line with previous work on identical fungal species (Zwinkels, van Oorschot, et al., 2025). In this study the EUC of pure basidiomycetous mycelia surpassed that of *R. microsporus* mycelium and their corresponding mushrooms, reaching up to 300 g MSG-eq/100 g DW. Thereby, the EUC of SSFF product reached half that of the highest EUC value of pure mycelium but outperformed all mushrooms. These results suggest that the EUC of optimized solid-state fermented products could surpass the umami intensity typically associated with mushrooms, highlighting the potential of basidiomycetes for creating appealing umami taste in food through solid-state fermentation.

Interestingly, the EUC of pure mycelium of a fungal species did not correspond with the EUC of their fermented product produced with that fungal species. Instead, the interaction between the fungal species and the substrate appears to be the key factor. This interaction suggests that high fungal proteolytic activity combined with a high protein content is likely required to achieve a high umami free amino acid content. Moreover, this interaction suggests that through substrate-species selection, the umami taste of these fermented products could be further optimized. Taken together, these findings underscore the potential of solid-state fermentation of plant-based food with basidiomycetes to offer a rich umami taste. This is a key factor in increasing the palatability and promoting the consumption of these sustainable protein sources.

Alkylpyrazines are important compounds in the aroma profile of roasted meat, mushrooms, and fermented foods like natto, and soy sauce (Moon et al., 2006; Rocchi et al., 2024; Zhang et al., 2018). Alkylpyrazines are nitrogenous cyclic compounds, which through alkylation greatly reduce their odour threshold, typically producing roasted, nutty, and meaty aromas (Supplementary table 1). They are generally formed in a Maillard reaction occurring at low water activity with amino acids (i.e. alanine, glycine, threonine, or lysine) and reducing sugars (i.e. glucose, fructose, and rhamnose) as reactants (Adams & Kimpe, 2009; Amrani-Hemaimi et al., 1995).

2,5-dimethypyrazine and tetramethylpyrazine were the most abundant compounds in raw, fermented BSG and lupin. Heating increased the abundance and/or diversity of the alkylpyrazines up to twelve compounds. This is in line with the alkylpyrazines found in pure mycelium of these species (Zwinkels, van Oorschot, et al., 2025). However, SSFF samples contained a higher diversity of alkylpyrazines in the raw product than pure mycelium did. Moreover, heated mycelium produced a higher diversity of alkylpyrazines than heated SSFF products. Furthermore, in pure mycelium, the highest abundance of alkylpyrazines was found in basidiomycetes, while in SSFF samples, alkylpyrazines were most abundant in *R. microsporus* fermented substrates. Interestingly, the composition of the substrate seems to have a large impact on the diversity and abundance of alkylpyrazines. Therefore, alkylpyrazine abundance in the mycelium is likely not a good predictor for alkylpyrazine formation in SSFF. Moreover, Sohail et al. (2022) and Zwinkels et al. (2025) reported that heated beef contains the highest diversity of alkylpyrazines, higher than chicken, mycelium, and mushrooms, with twelve pyrazine derivatives unique to this meat type. In line with these findings, several of these unique compounds were also detected in our SSFF products. Notably, 3-ethyl-2,5-dimethylpyrazine, identified by Sohail et al. (2022) as a distinctive aroma compound in chicken, beef, and pork, was present in the fermented samples.

The IVPDs of unfermented brown rice (82%) and lupin (92%) were slightly higher than those reported in literature for white rice (81%) and lupin (82%) (Villacrés & Rosell, 2021; Zwinkels et al., 2023). Contrarily, the IVPD of unfermented BSG (88%) was slightly lower than reported in barley (90-91%) (Zwinkels et al., 2023). The effect of SSFF on IVPD varies across the literature. Increasing IVPD have been reported in fermentations with *P. ostreatus, R. microsporus, Lentinula edodes*, and *Aspergillus oryzae* (Clark et al., 2022; Espinosa-Páez et al., 2017, 2021; Todorov et al., 2024; Villacrés & Rosell, 2021). However, fermentations with *Rhizopus* sp. were found to decrease the IVPD (da Silveira & Badiale-Furlong, 2009; Ranjan et al., 2019; Zwinkels et al., 2023). In this study, similar to the effect of fermentation on taste and aroma, the effect IVPD was dependent on substrate-species interaction. IVPD did not change significantly in brown rice, while digestibility decreased in BSG and increased in lupin fermented with *P. cinnabarinus*.

Although SSFF altered the IVPD significantly, the increased PDCAAS of fermented substrate were largely due to changes in the amino acid composition. Particularly the increase in lysine, in ten out of eleven fermentations, raised the PDCAAS. These results are in line with earlier SSFF by *R. microsporus* and *A. oryzae* on barley and white rice (Zwinkels et al., 2023). This increase in lysine is of particular importance because lysine is the main limiting amino acid in plant-based diets (Leinonen et al., 2019). Although basidiomycetes could increase the protein quality of specific substrates more than *R. microsporus*, fungal species from both phyla improve the protein quality of plant substrates. Nevertheless, as has been described in Zwinkels, Oorschot, et al. (2025), readily available carbon sources in the substrate (e.g. starch) could impact the protein content of the fermented product, by weight loss due to their consumption and conversion to CO_2_. Therefore, to assess the total quality of the protein source, both protein quality and protein content need to be evaluated.

For this reason, the utilizable amino acid content (UAA) was calculated, which evaluates both the protein quality as well as the total amino acid content (**Figure 5**). The total amino acid content was used instead of the crude protein content, because in fermented products, methods to detect nitrogen-based crude protein, may not accurately estimate protein content because deamination can increase non-amino nitrogen levels, leading to an overestimation of protein content (Zwinkels, Oorschot, et al., 2025). The change in UAA from unfermented substrates encompass changes arising from i) changes in protein quality, ii) changes in concentration of amino acids, due to loss of dry weight, and iii) loss of amino acids by deamination. This is illustrated by the following examples: i) fermentation of lupin with *P. cinnabarinus* increased the protein quality but slightly decreased total amino acids content (Supplementary table 2 & Supplementary figure 3); ii) fermentation of brown rice with *S. commune* increased total amino acid content with minimal change in protein quality; iii) fermentation of BSG with *P. cinnabarinus* decreased the total amino acid content, due to catabolism of amino acids, counteracting the increase in protein quality resulting in an overall decrease in UAA. These results indicate that both an increase in protein content and protein quality can be obtained with SSFF by basidiomycetes with the correct substrate-species formulation. A prerequisite for successful improvement of the nutritional quality appears to be the presence of a readily available carbon source. This prevents the utilization of amino acids as an alternative carbon source, while increasing the total amino acid content due to weight loss in form of non-protein compounds (e.g. carbohydrates). Similar observations were made by Stoffel et al. (2019), who showed that fermentation of BSG and grape bagasse by the basidiomycetes *Pleurotus albidus, Agaricus fuscosuccinea*, and *Agaricus blazei* had varying effects on the protein and amino acid content. They observed that when carbohydrates were metabolized, as seen with *P. albidus* in BSG, both protein and amino acid content increased. However, when carbohydrates were either absent (in grape bagasse) or could not be utilized (by *Agaricus* species), total amino acid content decreased, even though crude protein content still increased.

Finally, basidiomycetous species improved the total UAA content of the cereal (brown rice) and the legume (lupin) more effectively than *R. microsporus*, by increasing the AAS of lysine and the total concentration of amino acids through a loss of dry weight. This is particularly relevant since cereals currently provide approximately 30% of the protein in human diets, with maize, wheat, and rice as the dominant crops (FAO, 2024). This ratio is expected to rise with an increase in plant-based diets (Leinonen et al., 2019). However, cereals typically have a high calorie-to-protein ratio and are deficient in lysine (Sá & House, 2024). Therefore, the combined effect of increasing the PDCAAS of lysine and elevation of the total amino acid content, due to removal of non-nitrogen caloric nutrients, significantly enhances the potential of cereals to contribute to meeting dietary protein requirements.

In conclusion, solid-state fermentation with edible basidiomycetes could increase in the utilizable amino acid content as well as the umami taste, particularly in *P. cinnabarinus* fermentations. These results imply that solid-state fermentation with edible basidiomycetes could produce novel alternative protein-enriched foods or ingredients with improved nutritional value and umami taste, which are important drivers for alternative protein adoption (Onwezen et al., 2021; Schmidt & Mouritsen, 2022). Furthermore, the altered texture, defined by the transformation from loose kernels to a solid patty, transforms the perception of the food. Thereby, these novel cereal-based tempeh-like products could better serve as the protein source of the meal. However, more refinements in substrate-species formulation are required to optimize the fermented food product in terms of protein quality, taste, and texture. Finally, legislative hurdles might have to be taken as such products likely fall under the EU Novel Food regulation (2015/2283) (Molitorisová et al., 2025).

## Supporting information

Supplementary file Zwinkels et al. 2025

## CRediT authorship contribution statement

**Jasper Zwinkels:** Conceptualization, Methodology, Investigation, Formal analysis, Data curation, Validation, Writing – original draft. **Oscar van Mastrigt:** Conceptualization, Methodology, Writing – review & editing, Supervision. **Eddy J. Smid:** Conceptualization, Methodology, Writing – review & editing, Supervision, Funding acquisition, Project administration.

## Acknowledgements

We would like to thank Stef van Oorschot for his contribution to the production and analysis of solid-state fermented BSG. Additionally, the exploratory studies of Philip van Koolwijk, Ran Xu, and Alessia de Chaud were very valuable for this paper. We gratefully acknowledge Dr. Rebecca Rocchi for her inspiration in the graphical design. Lastly, we would like to thank Dr. Nikkie van der Wielen, Michel Breuer and Dr. Leon de Jonge for their advice and performing the amino acid extraction and analysis.

## Data availability

Data will be made available on request.

