## Supplementary file Zwinkels et al. 2025 for "Solid-state fermentation with mushroom mycelium elevates plant protein quality and umami taste"

### 1 Supplementary material

#### 1.1 Supplementary tables

**Supplementary Table 1.** Odor description and odour threshold in water (µg/L) of alkylpyrazines present in mycelia, fruiting bodies or meat (Cherniienko et al., 2022; Shi et al., 2025; Sohail et al., 2022).

| **Compounds** | **Odor description** | **Odor threshold in water (in µg/L)** |
| --- | --- | --- |
| 2,3,5-Trimethyl-6-ethylpyrazine | chocolate, cocoa, coffee, sweet, hazelnut, roasted | 0,002-0,024 |
| 2,3-Dimethylpyrazine | nutty, cocoa, peanut butter, coffee, caramel, roasted potato, musty | 400-2500  15 mg/kg |
| 2,5-Dimethylpyrazine | cocoa, roasted, nutty, beef, woody, grassy, medicinal, earthy | 1700-2600 |
| 2,6-Dimethylpyrazine | ether, cocoa, nutty, roasted, beefy, coffee, buttermilk | 400-9000 |
| 2-Acetyl-3-methylpyrazine | nutty, popcorn, roasted corn, dirt, burnt, sweet | 62 |
| 2-Ethyl-3,5-dimethylpyrazine | burnt, coffee, nutty, roasted, woody, potato-chip-like, | 0,04-2 |
| 2-Ethyl-3-methylpyrazine | nutty, musty, corn, raw, earthy, bready | 130-500 |
| 2-Ethyl-6-methylpyrazine | roasted, potato | 40 |
| 2-Methyl-5-(1-propenyl)-(E)-pyrazine | Roasted | - |
| 2-Methyl-5-(1-propenyl)-(Z)-pyrazine | Roasted | - |
| 3,5-Diethyl-2-methylpyrazine | nutty, meaty, vegetable | 0,9 (in diethyl ether) |
| 3,5-Dimethyl-2-isobutylpyrazine | roasted, sweet, green | - |
| 3-Ethyl-2,5-dimethylpyrazine | nutty, roasted sunflower seeds | 35 |
| 6,7-Dihydro-2,5-dimethyl-5H-cyclopentapyrazine | - | - |
| Methylpyrazine | pungent, sweet, corn-like, nutty, chocolate, hazelnut, green | 27.000-100.000 |
| *Tetramethylpyrazine* | nutty, musty, chocolate, coffee, cocoa, burnt, musty, vanilla | 1000-38.000 |
| Trimethylpyrazine | nutty, earthy, powdery, cocoa, potato, roasted | 90-1800 |
| 2-Isoamyl-6-methylpyrazine | Green, earthy, roasted | - |
| Ethylpyrazine | Nutty, peanut | 0.2-1 |
| 2,3-Dimethyl-5-isopentylpyrazine | Earthy, roasted | 6000 |
| 1-Methylethenylpyrazine | Spicy, roasted, vegetable | - |

**Supplementary Table 2.** Amino acid content (mg/g DW), essential amino acid percentage (%), and total amino acid content (mg/g DW) of unfermented and fermented brown rice, brewer’s spent grain (BSG), and lupin.

|  | **Brown rice** | | | | | | | | | | | | **BSG** | | | | | | | | | | **Lupin** | | | | | | |
| --- | --- | --- | --- | --- | --- | --- | --- | --- | --- | --- | --- | --- | --- | --- | --- | --- | --- | --- | --- | --- | --- | --- | --- | --- | --- | --- | --- | --- | --- |
| Amino acid | **Unfermented** | | ***S. commune*** | | ***V. volvacea*** | | ***P. cinnabarinus*** | | ***P. pulmonarius*** | | ***R. microsporus*** | | **Unfermented** | | ***S. rugosoannulata*** | | ***P. cinnabarinus*** | | ***P. pulmonarius*** | | ***R. microsporus*** | | | **Unfermented** | | ***P. cinnabarinus*** | | ***R. microsporus*** | |
| (mg/g DW) | **mean** | **sd** | **mean** | **sd** | **mean** | **sd** | **mean** | **sd** | **mean** | **sd** | **mean** | **sd** | **mean** | **sd** | **mean** | **sd** | **mean** | **sd** | **mean** | **sd** | **mean** | **sd** | | **mean** | **sd** | **mean** | **sd** | **mean** | **sd** |
| Alanine | 5.99 | 0.03 | 8.31 | 0.30 | 6.45 | 0.14 | 7.37 | 0.42 | 6.39 | 0.14 | 6.99 | 0.13 | 10.40 | 0.05 | 9.61 | 1.03 | 6.44 | 0.05 | 9.63 | 0.69 | 10.61 | 0.21 | | 14.29 | 0.02 | 16.17 | 0.15 | 15.78 | 0.65 |
| Arginine | 8.62 | 0.02 | 11.11 | 0.15 | 7.51 | 0.14 | 8.19 | 0.35 | 8.00 | 0.41 | 8.64 | 0.13 | 11.86 | 0.28 | 11.81 | 2.88 | 5.02 | 0.14 | 8.06 | 0.88 | 11.06 | 0.26 | | 39.51 | 0.78 | 31.48 | 0.46 | 31.77 | 0.08 |
| Aspartic acid | 9.31 | 0.08 | 12.20 | 0.32 | 10.21 | 0.30 | 10.51 | 0.40 | 9.35 | 0.23 | 9.91 | 0.13 | 15.06 | 0.11 | 15.14 | 1.85 | 10.14 | 0.12 | 13.55 | 1.23 | 16.68 | 0.45 | | 43.91 | 0.29 | 43.95 | 0.20 | 41.37 | 0.31 |
| Cysteine | 2.27 | 0.02 | 3.00 | 0.07 | 2.45 | 0.01 | 2.73 | 0.08 | 2.22 | 0.02 | 2.81 | 0.02 | 4.22 | 0.06 | 4.20 | 0.03 | 3.03 | 0.04 | 3.39 | 0.27 | 4.48 | 0.08 | | 6.28 | 0.08 | 7.14 | 0.08 | 5.70 | 0.09 |
| Glutamic acid | 17.91 | 0.18 | 23.93 | 0.56 | 17.43 | 0.14 | 18.99 | 0.61 | 17.15 | 0.24 | 18.01 | 0.12 | 40.49 | 0.76 | 33.74 | 5.40 | 17.39 | 1.09 | 25.82 | 0.65 | 38.08 | 1.79 | | 80.00 | 0.85 | 60.65 | 0.53 | 70.05 | 0.62 |
| Glycine | 4.89 | 0.03 | 6.43 | 0.18 | 5.33 | 0.14 | 5.71 | 0.24 | 4.92 | 0.13 | 5.00 | 0.03 | 9.10 | 0.05 | 8.53 | 0.98 | 6.07 | 0.05 | 7.92 | 0.70 | 9.12 | 0.22 | | 16.59 | 0.06 | 18.75 | 0.21 | 15.84 | 0.06 |
| Histidine | 2.57 | 0.05 | 3.44 | 0.07 | 2.63 | 0.07 | 3.00 | 0.14 | 2.52 | 0.03 | 2.94 | 0.04 | 4.83 | 0.14 | 4.67 | 0.53 | 2.57 | 0.05 | 3.83 | 0.17 | 5.04 | 0.14 | | 9.87 | 0.11 | 10.63 | 0.13 | 10.36 | 0.15 |
| Isoleucine | 4.43 | 0.05 | 6.10 | 0.19 | 5.08 | 0.13 | 5.25 | 0.25 | 4.89 | 0.14 | 4.72 | 0.01 | 8.80 | 0.12 | 7.90 | 0.67 | 3.80 | 0.09 | 6.47 | 0.39 | 9.15 | 0.26 | | 19.40 | 0.14 | 17.99 | 0.23 | 19.09 | 0.16 |
| Leucine | 8.51 | 0.07 | 11.26 | 0.30 | 9.27 | 0.24 | 9.51 | 0.41 | 8.93 | 0.21 | 8.72 | 0.03 | 16.33 | 0.23 | 14.13 | 0.76 | 6.50 | 0.09 | 11.34 | 0.93 | 16.06 | 0.45 | | 33.26 | 0.17 | 26.92 | 0.30 | 31.37 | 0.13 |
| Lysine | 3.98 | 0.02 | 5.32 | 0.13 | 4.85 | 0.22 | 5.28 | 0.29 | 4.62 | 0.17 | 4.71 | 0.10 | 9.46 | 0.22 | 10.05 | 0.79 | 6.93 | 0.57 | 6.95 | 0.13 | 10.29 | 0.27 | | 19.61 | 0.26 | 19.42 | 0.24 | 20.67 | 0.17 |
| Methionine | 2.95 | 0.03 | 3.65 | 0.09 | 2.93 | 0.05 | 3.08 | 0.11 | 2.87 | 0.07 | 3.07 | 0.05 | 4.15 | 0.05 | 3.56 | 0.11 | 2.74 | 0.21 | 2.75 | 0.22 | 4.06 | 0.11 | | 3.32 | 0.01 | 3.77 | 0.09 | 3.56 | 0.01 |
| Proline | 4.85 | 0.04 | 6.35 | 0.16 | 5.31 | 0.12 | 5.54 | 0.22 | 5.06 | 0.14 | 4.89 | 0.05 | 19.17 | 0.46 | 13.32 | 1.45 | 5.54 | 0.10 | 9.58 | 1.25 | 16.73 | 0.94 | | 16.98 | 0.10 | 16.74 | 0.20 | 16.59 | 0.16 |
| Serine | 5.34 | 0.10 | 7.18 | 0.20 | 5.77 | 0.13 | 5.95 | 0.22 | 5.55 | 0.12 | 5.52 | 0.04 | 9.62 | 0.12 | 8.68 | 1.45 | 5.12 | 0.17 | 6.98 | 0.39 | 9.49 | 0.24 | | 22.20 | 0.13 | 19.01 | 0.23 | 19.73 | 0.30 |
| Threonine | 3.81 | 0.04 | 5.31 | 0.18 | 4.88 | 0.15 | 4.97 | 0.23 | 4.70 | 0.17 | 4.35 | 0.02 | 8.28 | 0.08 | 7.77 | 1.04 | 4.68 | 0.03 | 7.25 | 0.45 | 8.48 | 0.14 | | 16.36 | 0.07 | 17.21 | 0.19 | 16.55 | 0.05 |
| Tryptophan | 1.23 | 0.00 | 1.63 | 0.06 | 1.46 | 0.00 | 1.51 | 0.06 | 1.33 | 0.03 | 1.32 | 0.03 | 2.59 | 0.07 | 1.83 | 0.15 | 2.80 | 0.20 | 1.86 | 0.07 | 2.68 | 0.02 | | 2.89 | 0.03 | 3.71 | 0.04 | 2.94 | 0.06 |
| Valine | 6.19 | 0.06 | 8.28 | 0.24 | 6.45 | 0.18 | 6.89 | 0.30 | 6.26 | 0.12 | 6.44 | 0.09 | 11.90 | 0.10 | 11.43 | 0.81 | 6.11 | 0.88 | 9.39 | 1.13 | 11.71 | 0.29 | | 17.80 | 0.07 | 18.01 | 0.21 | 17.89 | 0.21 |
| Essential amino acids (%) | 38.71 | 0.03 | 38.86 | 0.07 | 40.82 | 0.22 | 40.41 | 0.12 | 40.45 | 0.21 | 39.86 | 0.06 | 37.88 | 0.10 | 39.58 | 2.65 | 41.27 | 1.90 | 39.51 | 0.35 | 39.17 | 0.30 | | 35.55 | 0.13 | 37.64 | 0.05 | 37.77 | 0.13 |
| Total amino acids | 92.8 | 0.59 | 123.5 | 3.18 | 98.0 | 2.07 | 104.5 | 4.21 | 94.8 | 2.09 | 98.1 | 0.71 | 186.3 | 2.59 | 166.4 | 17.78 | 94.9 | 0.73 | 134.8 | 8.29 | 183.7 | 5.80 | | 362.3 | 2.61 | 331.6 | 3.13 | 339.3 | 1.55 |

#### 1.2 Supplementary figures

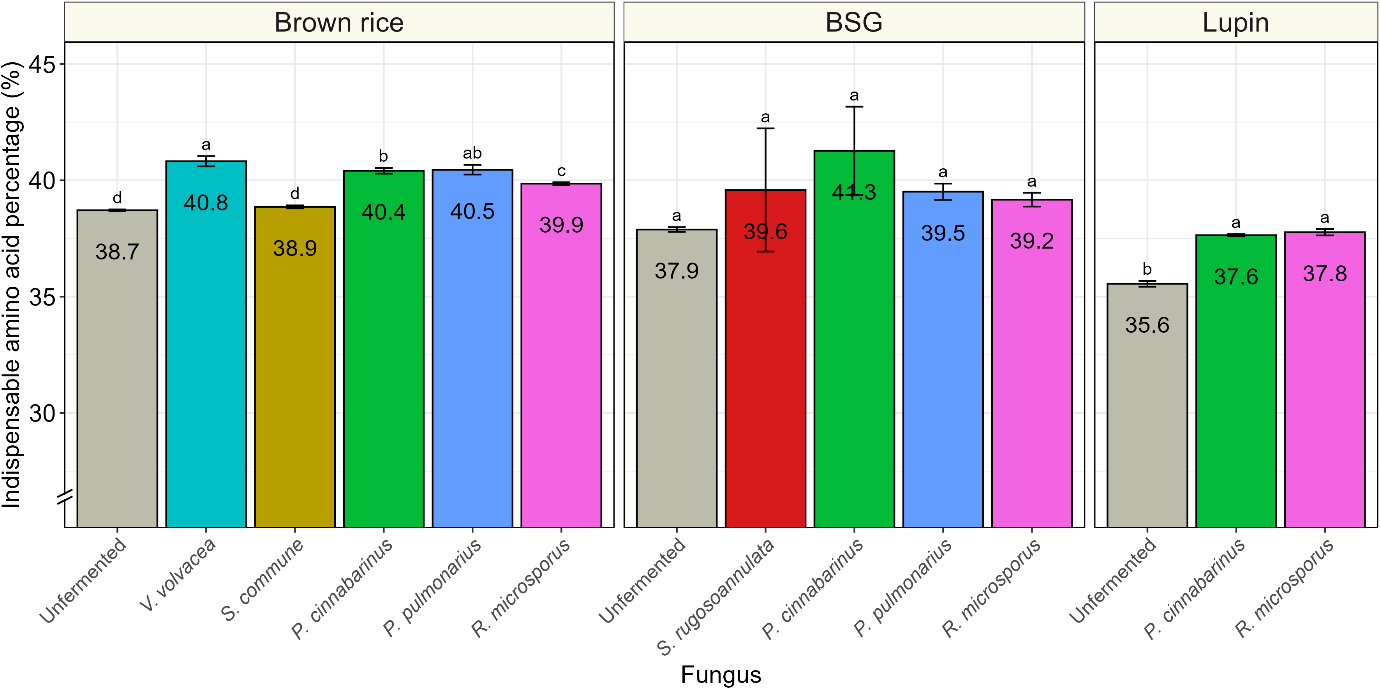

**Supplementary figure 1.** Essential amino acid percentage (% of total amino acids) in unfermented and fermented substrates.

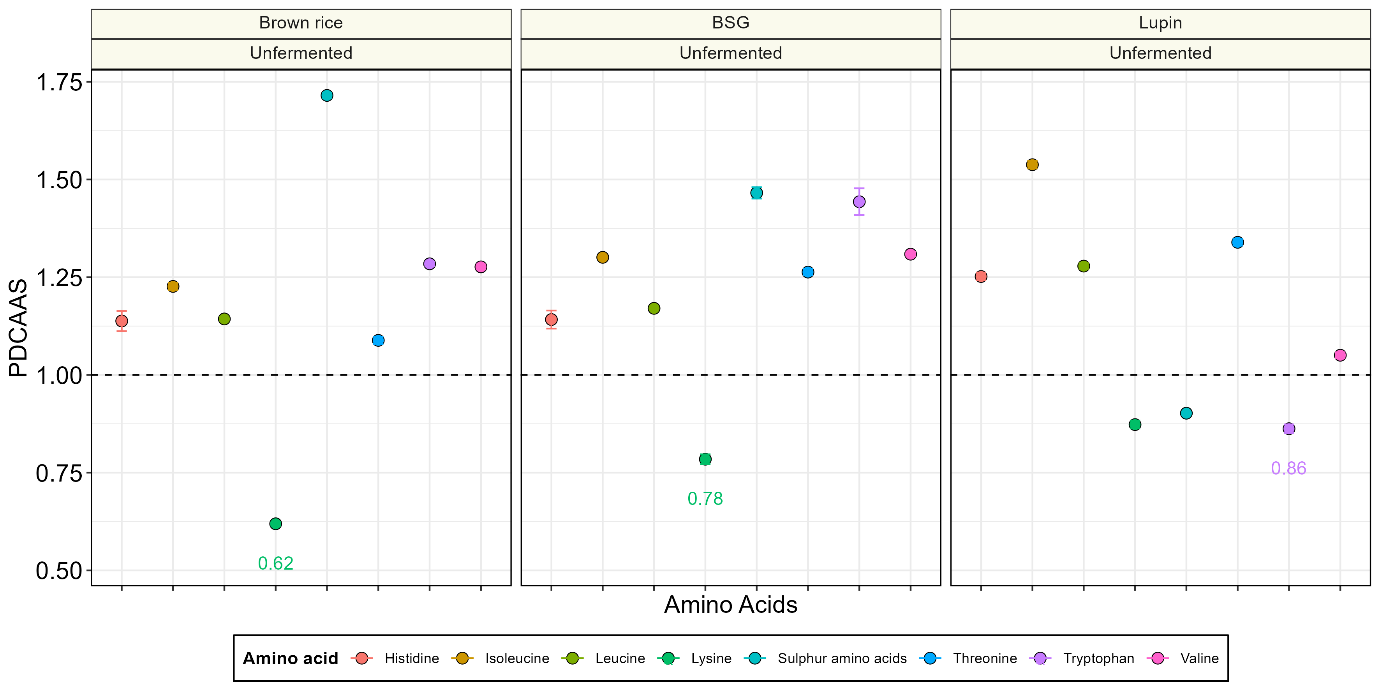

**Supplementary figure 2.** Protein digestibility corrected amino acid score (PDCAAS) of unfermented brown rice, BSG and lupin. Colours indicate indispensable amino acid. The PDCAAS for the limiting amino acid is provided in text. Aromatic amino acids (phenylalanine and tyrosine) were not measured.

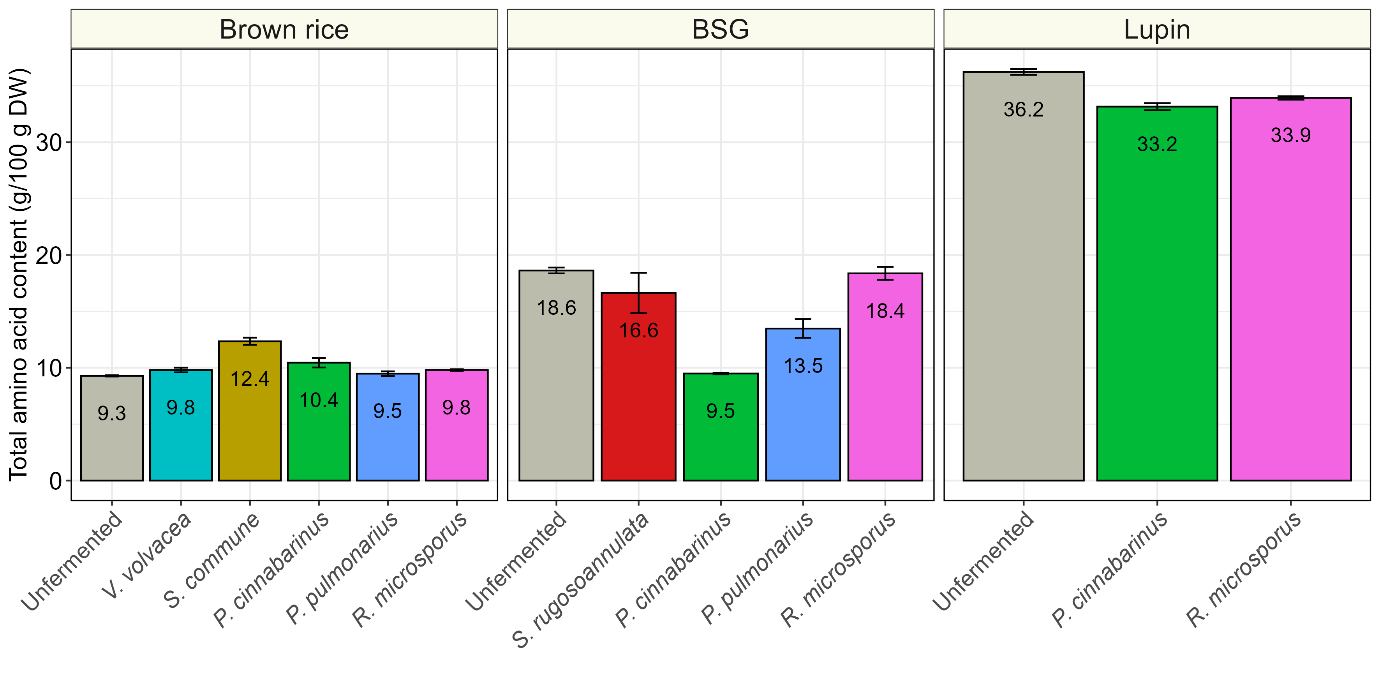

**Supplementary figure 3.** Total amino acid content (g/100 g DW) of unfermented and fermented substrates.
